# Bi-functional particles for real-time acidification and proteolysis multiplex assay in macrophages

**DOI:** 10.1101/2023.04.03.535364

**Authors:** Alba Méndez-Alejandre, Benjamin Bernard Armando Raymond, Matthias Trost, José Luis Marín-Rubio

## Abstract

Phagosome acidification and proteolysis are essential processes in the immune response to contain and eliminate pathogens. In recent years, there has been an increased desire for a rapid and accurate method of assessing these processes in real-time. Here, we outline the development of a multiplexed assay that allows simultaneous monitoring of phagosome acidification and proteolysis in the same sample using silica beads conjugated to pHrodo and DQ BSA. We describe in detail how to prepare the bi-functional particles and show proof of concept using differentially activated macrophages. This multiplexed spectrophotometric assay allows rapid and accurate assessment of phagosome acidification and proteolysis in real-time and could provide valuable information for understanding the immune response to pathogen invasion.

## INTRODUCTION

Phagocytosis is an essential process for the innate immune system, as it helps to protect the body from pathogens and other harmful substances. Additionally, this process is essential for tissue repair, to regulate the inflammatory response and to present antigens to other immune cells. In this process, cells engulf and absorb foreign particles larger than 0.5 μm in diameter, such as microorganisms, foreign particles, or cellular debris (Kinchen & Ravichandran, 2008). While many types of cells are able to phagocytose, this is largely performed by professional phagocytes such as macrophages, neutrophils, monocytes, dendritic cells, and osteoclasts. Macrophages are the most important type of phagocytes, as they are able to recognize and engulf foreign particles, as well as release cytokines that help to activate other cells in the immune response (Uribe-Querol & Rosales, 2020). After phagocytic uptake, the newly formed phagosome undergoes a series of changes in protein and membrane composition, termed phagosome maturation (Garin et al, 2001; Trost et al, 2009; Pauwels et al, 2017; Hipolito et al, 2019). The process begins when a cell binds to a particle and forms a phagocytic cup around it, then uses its pseudopods to engulf the particle, allowing it to be pulled into the cell. In the early stages, the phagosome membrane fuses with endocytic vesicle membranes that contain important factors such as the v-ATPases, which pump protons (H^+^) from the cytosol and acidify the lumen of the phagosome (reaching pH∼5), and the major histocompatibility complex (MHC) proteins, which are essential for antigen presentation (Ackerman et al, 2003; Pauwels et al, 2017). The phagosome also increases in size, allowing it to accommodate more material and absorb more substances (Hipolito et al, 2019). Finally, the phagosome fuses with the lysosome forming the phagolysosome, which delivers hydrolytic enzymes such as proteases, nucleases, glycosidases, lipases, and reactive oxygen species (ROS) to kill and digest the pathogens (Garin et al, 2001; Xu & Ren, 2015). In this stage, proteases can only be activated under acidic conditions to digest the pathogen and process the antigen for its presentation (Hipolito et al, 2019). By exposing certain antigens on its outer surface, the phagosome can also activate the immune system and help the body effectively fight off foreign invaders. It is therefore an essential process for the survival and protection of the cell.

The acidification and proteolysis of phagosomes in phagocytic cells are key processes that have important implications for cellular homeostasis. When phagocytic cells are unable to properly acidify and proteolyse phagosomes, it can lead to alterations in the immune response to infections and pathogens, which can have adverse clinical consequences (Engelich et al, 2001; Andrews & Sullivan, 2003; Uribe-Querol & Rosales, 2017). In order to better understand these processes, a variety of techniques have been implemented to monitor single processes, including immunoblotting, quantitative PCR, and flow cytometry to measure changes in the levels of proteolytic enzymes, changes in the mRNA expression of proteolytic enzymes or changes in phagosome formation and determine the acidification state of the phagosome (Russell et al, 2009; Sanman et al, 2016; Frost et al, 2017). Additionally, other techniques have been used to visualize the uptake of fluorescently-labelled particles into phagosomes and track changes in their acidification and maturation. These assays rely on pH-sensitive fluorescent dyes, particles bound to *E. coli*, bioparticles labelled with pH-sensitive and pH-insensitive fluorochromes (Yates et al, 2005; Yates & Russell, 2008; Russell et al, 2009; Lee et al, 2021). However, none of these mehtods allows for the investigation of both acidifcation and proteolysis in parellel.

To address this need, we report here a detailed method to prepare bi-functional particles for the simultaneous quantification of pH and proteolytic activity inside phagosomes during maturation using a spectrophotometer. For this, we coated silica beads with pHrodo and DQ BSA to measure the acidity and hydrolysis activity in phagocytic cells.

## MATERIALS AND REAGENTS

### Materials

- 50 mg/mL, 3.0 µm carboxylate-modified silica particles (Kisker Biotech)
- 96-well black clear bottom polystyrene microplates (Corning Life Sciences)
- Spectrophotometer, SpectraMax Gemini EM (Molecular Devices).

### Primary macrophages and macrophage cell line

- Bone marrow differentiated macrophages (BMDMs) (Heap et al, 2021) from 10 to 12-week-old C57BL/6NTac mice (UKRI-MRC Harwell). Newcastle University ethical committee approved animal work and manipulation was performed under UK Home Office project licence.
- RAW 264.7 (ATCC, TIB-71) cells. ATCC routinely performs cell line authentication, using short tandem repeat profiling as a procedure. Cell experimentation was always performed within a period not exceeding 6 months after resuscitation in mycoplasma-free culture conditions.

### Reagents and buffers

- Dubecco’s modified eagle medium (DMEM)
- Iscove’s Modified Dulbecco’s Media (IMDM)
- 100 U/ml penicillin-streptomycin
- Foetal bovine serum (FBS)
- 2 mM L-Glutamine
- Interferon gamma (IFN-γ) (PeproTech)
- Interleukin 4 (IL-4) (PeproTech)
- Binding buffer: 1 mM CaCl2, 2.7 mM KCl, 0.5 mM MgCl2, 5 mM dextrose, 10 mM HEPES and 10 % FBS in PBS pH 7.2
- Sodium borate buffer: 100 mM boric acid in purified water, pH 8.0
- Glycine buffer: 250 mM glycine in PBS pH 7.2
- Cyanamide buffer: 25 mg/mL in purified water, make fresh
- Sodium azide 1 %
- 5 mg/mL Alexa Fluor 405 (AF405) carboxylic acid, succinimidyl ester (Invitrogen) in DMSO
- 2 mg/mL, DQ red BSA (Invitrogen) in sodium borate buffer
- 5 mg/mL pHrodo green, succinimidyl ester (Invitrogen) in DMSO
- 100 µM bafilomycin A1 (Sigma-Aldrich)

## METHODS

### Preparation of bi-functional particles (Figure 1. 1–5)

1. Take 100 µL of 3.0 µm carboxylate-modified silica beads.

**Figure 1.**
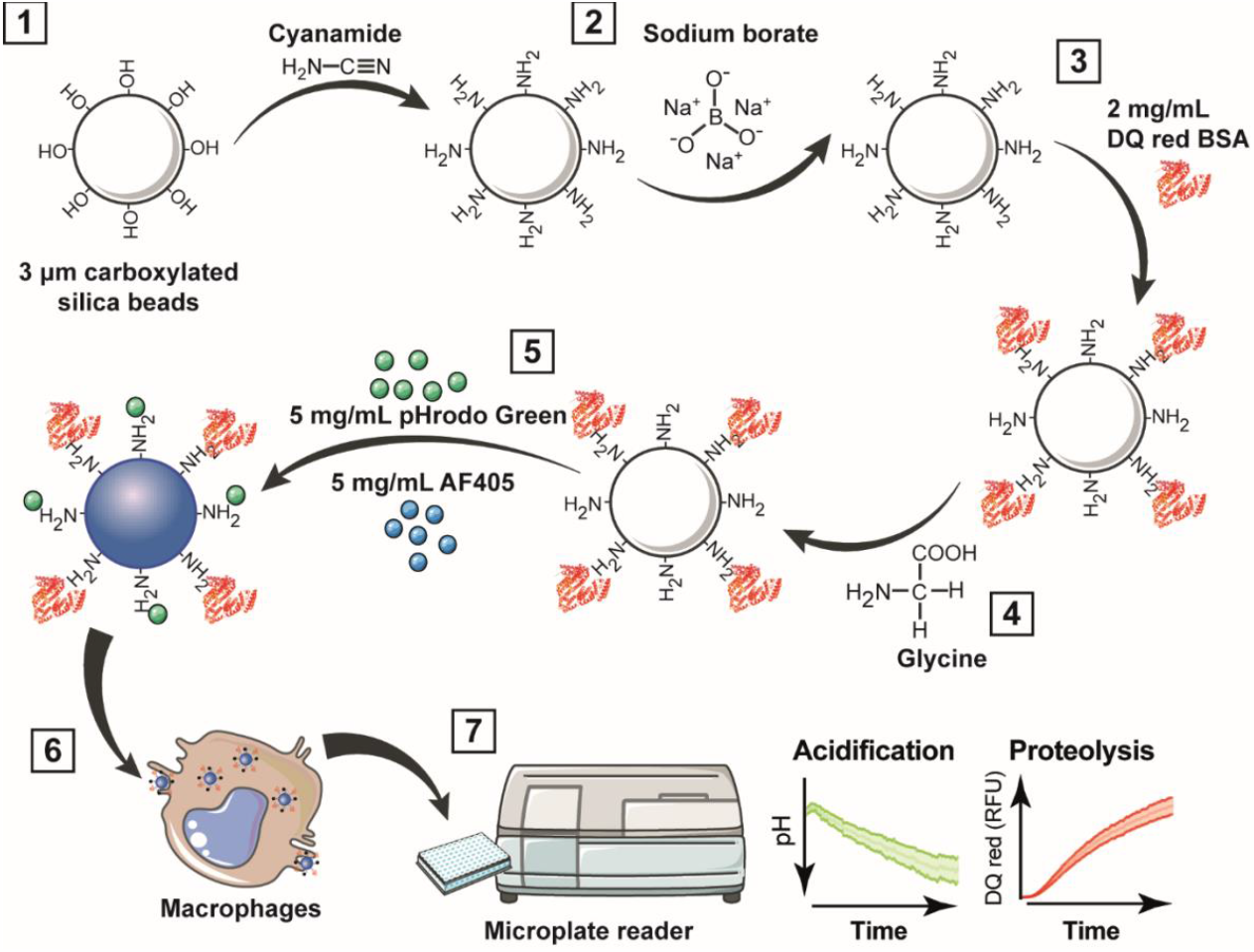
Workflow for multiplexing analysis of phagosome acidification and proteolysis. **1)** Carboxyl functionalized silica nanoparticles or micro beads are wash with the heterobifunctional crosslinker cyanamide; **2)** excess amines are removed by washing with sodium borate; then, **3)** beads are incubated overnight with 2 mg/mL DQ red BSA in sodium borate buffer. **4)** After the incubation, the beads are washed with glycine to quench unreacted cyanamide; and **5)** incubated with 5 mg/mL pHrodo green and 5 mg/mL AF405 in sodium borate buffer for 1 h. **6)** Bi-functional particles are added to macrophages, and **7)** acidification and proteolysis are measured in real time simultaneously using a fluorescence microplate reader. RFU, relative fluorescence units.
2. Wash the beads three times with 1 mL PBS at 2,000 x g for 1 min using a benchtop centrifuge.
3. Incubate the beads with 500 µL of the heterobifunctional crosslinker cyanamide (cyanamide buffer) at 900 rpm for 15 min at room temperature.
4. Wash twice with 1 mL of sodium borate buffer at 2,000 x g for 3 min to remove soluble amine groups.
5. Incubate the beads with 500 µL of 2 mg/mL DQ red BSA in sodium borate buffer for 18 h under constant agitation (900 rpm) at RT in the dark.
6. After the incubation, wash the beads twice with glycine buffer at 2,000 x g for 2 min to quench unreacted cyanamide.
7. Wash the beads twice with sodium borate buffer at 2,000 x g for 3 min to remove soluble amine groups.
8. Incubate the beads with 1 mL of 5 mg/mL pHrodo green and 5 mg/mL AF405 for 1 h in sodium borate buffer under constant agitation (900 rpm) at room temperature in the dark.
9. Wash the beads twice in PBS at 2,000 x g for 3 min.
10. Resuspend in 500 µL PBS with 1 % sodium azide and stored in low-binding tubes at 4°C within a period of not more than one month.

### Fluorogenic kinetic assay in live cells (Figure 1. 6)

1. BMDMs or RAW 264.7 cells were seeded on 96-well black clear bottom polystyrene microplates at 100,000 viable cells per well.
2. Treat cells with 20 μg/ml interferon gamma (IFN-γ) or 20 μg/ml interleukin 4 (IL-4) for 24 h and 48 h before the assay.
3. Removed medium and wash cells twice with warm Binding buffer.
4. Add 100 μM bafilomycin A1 for 1 h before the assay as a v-ATPase inhibitor of phagosome maturation (negative control). Maintain the treatment for the rest of the experiment.
5. Add bi-functional particles at a dilution of 1:200 in Binding buffer and incubate for 10 min at 37°C in the incubator to allow the particles attach to the macrophages.
6. Wash the cells twice with Binding buffer.

#### Note

To validate the uptake of these bi-functional particles inside the cells, RAW

264.7 cells were imaged on a Zeiss LSM800 at 40X (oil immersion) using Zen software with the following lasers set to 1 % power: 405 nm, 488 nm, and 561 nm. The electronically switchable illumination and detection module (ESID) was also used to collect transmitted light (**Figure S1**).

7 Read the plate immediately with the microplate reader.

### Spectrofluorometer setup and operation (Figure 1. 7)

1. Kinetic readings were collected over time with multiple readings during 6 h with intervals of 4 min at maximal readings per well at 37°C.
2. Multiple-wavelength readings (emission/excitation) were performed at the same time. Optimal measurements were taken using the desired wavelengths as follows:

- Em/Ex 585/625 nm for DQ red BSA
- Em/Ex 395/435 nm for AF405
- Em/Ex 490/530 nm for pHrodo green

### Assessment of phagosomal pH

The average of three technical replicates were calculated and blank values were subtracted from each data set. The relative fluorescent units (RFUs) were calculated by normalising pHrodo green dye intensity against the calibration Alexa Fluor 405 intensity. To calculate the pH, a cubic polynomial regression curve (*f p(x): ax*^*3*^ *+ bx*^*2*^ *+ cx + d*) was analysed by obtaining measurements of the coupled particles at a known pH range (7.5 – 4.0) in each experiment to assess precision. The RFUs were then interpolated to this pH curve to obtain the acidification results (**Figure 2A** and **Figure S2A**). Excitation/emission corresponding to DQ red BSA (585/625 nm) was not affected by pH (**Figure S3A**).

**Figure 2.**
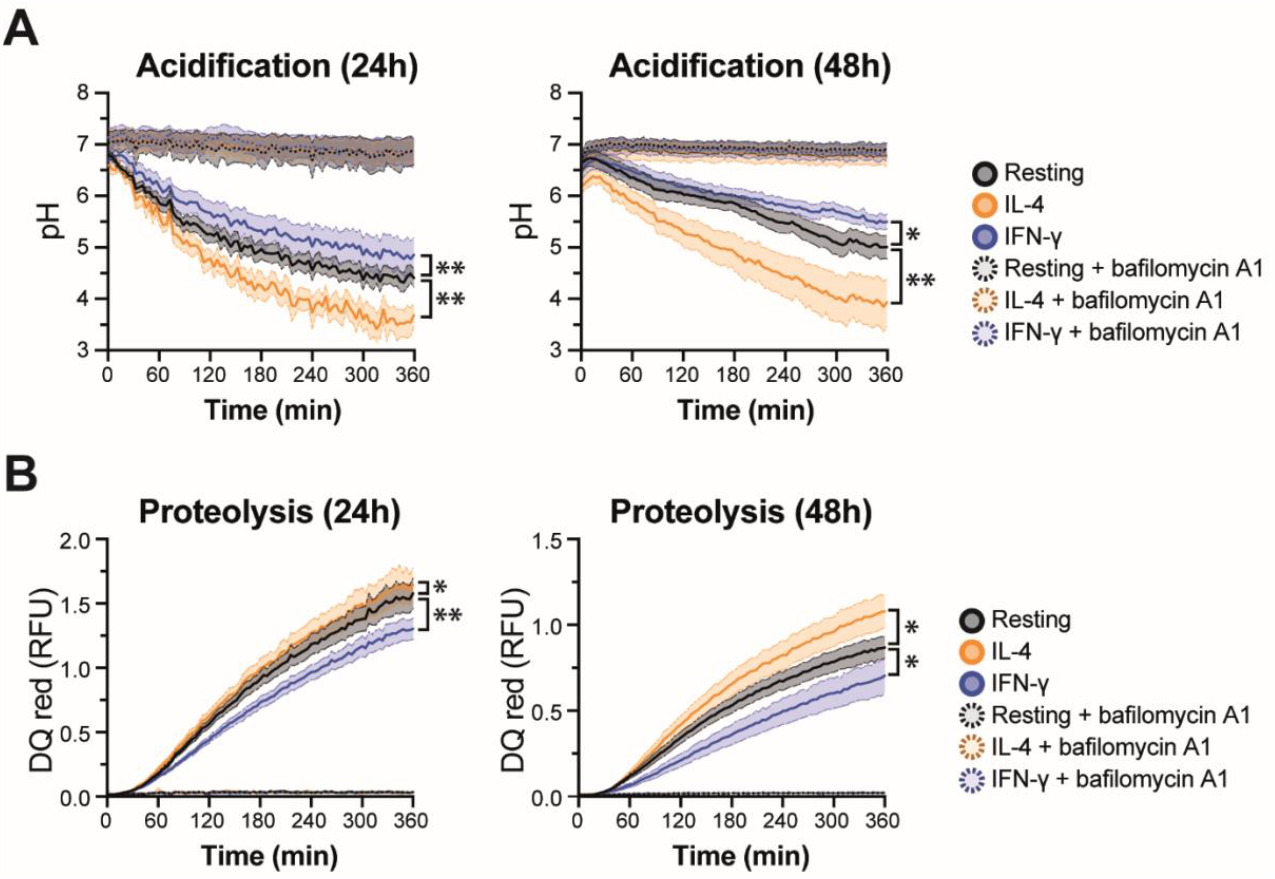
Real-time multiplex phagosome acidification and proteolysis analysis in BMDMs. **A)** Acidification and **B)** proteolysis were measured in BMDMs untreated (resting) or treated with 20 μg/mL IFN-γ or 20 μg/mL IL-4 for 24 h (left panels) and 48 h (right panels). Bafilomycin A1 was used as a negative control of phagosome maturation. Friedman one-way ANOVA test followed by Dunn post hoc test. The statistical significance of the comparisons with resting is indicated as follows: **, P ≤ 0.01; *, P ≤ 0.05. Error bars represent six biological replicates. RFU, relative fluorescence units.

### Assessment of phagosomal proteolysis

To measure the phagosomal proteolysis the same approach for pHrodo green was followed with DQ red BSA. In short, blank values were subtracted from each data set. The relative fluorescent units (RFUs) were calculated by normalising DQ red BSA dye intensity against the calibration Alexa Fluor 405 intensity (**Figure 1B** and **Figure S2B**). To validate the proteolysis of DQ red BSA, the particles were incubated at 37°C with 1 μg/μL trypsin TPCK (Worthington-Biochem) for 2 h to induce the proteolysis of BSA and measure in the spectrofluorometer. Excitation/emission corresponding to pHrodo Green (490/530 nm) was not affected by trypsin (**Figure S3B**).

### Statistical analysis

Statistical analyses were performed in GraphPad Prism (version 9.3.1). A nonparametric test was used for paired data: Friedman one-way ANOVA test followed by Dunn post hoc test; p-value < 0.05 was considered statistically significant.

## DISCUSSION

We described in detail and developed a method to prepare a bi-functional particle probe to simultaneously measure acidification and proteolysis during phagosome maturation in macrophages. This assay measures in real time the gradual increase in pH and acidic hydrolases during phagosomal acidification and proteolysis, respectively. Generally, compared to existing techniques for measuring chemical reactions within living cells, the pH of the phagosome is measured with pH-sensitive pHrodo dye or carboxyflorescein, attached to microscopic beads, which will emit higher fluorescence when exposed at acidic pHs (Yates et al, 2005; Sanman et al, 2016; Frost et al, 2017; Bilkei-Gorzo et al, 2022). However, pHrodo-labelled bioparticles showed a more precise and efficient quantification of phagosomal pH, under different assays, such as spectrofluorometry and flow cytometry (Colas et al, 2014). Our assay combines multiple fluorescent dyes to measure the pH and protein content inside the phagosome in real-time. For proteolysis, DQ red BSA was used. The close proximity of the dye molecules on BSA quenches the fluorescence while BSA is intact and when digested by acidic hydrolases found in the phagolysosome, its fluorescence increases when DQ red containing peptides are released (Frost et al, 2017). This combination of fluorescent dyes provides a powerful tool for the simultaneous monitoring of two important processes in a single sample.

The assay is able to accurately assess changes in both phagosome acidification and proteolysis and can be adapted for use in a variety of cell types. Furthermore, the assay is designed to be robust and easily adapted to be high throughput. This new approach to multiplexing will enable researchers to rapidly and accurately assess the effects of external stimuli on phagosome acidification and proteolysis in a wide range of cells and organisms. Lastly, because this method can be applied to labelled bacteria or apoptotic/necroptotic cells, this technique will be broadly applicable to determine the mechanisms of action of pathogens cells that evade host immunity by manipulating phagosome functions.

## Abbreviations

AF405: Alexa Fluor 405
BMDM: bone marrow-derived macrophages
DQ BSA: dye-quenched bovine serum albumin
IFN-γ: interferon-gamma
IL-4: interleukin-4
RFU: relative fluorescence units.

## ASSOCIATED CONTENT

### Supporting information

Figure S1. Bi-functional particles in RAW 264.7 cells.

Figure S2. Real-time multiplex phagosome acidification and proteolysis analysis in RAW 264.7 cells.

Figure S3. Quality control of bi-functional particles.

## AUTHOR INFORMATION

## Authorship contributions

A.M.-A. and J.L.M.-R performed and analysed most experiments. B.B.A.R. performed additional experiments. J.L.M.-R., A.M.-A., and M.T. wrote the manuscript. M.T. and J.L.M.-R. conceived the original idea and provided supervision. All authors provided critical feedback and helped shape the research, analysis, and manuscript.

## Notes

The authors declare no competing financial interest.

## ACKNOWLEDGEMENTS

This research was funded by a Wellcome Trust Investigator Award (215542/Z/19/Z) to M.T. J.L.M-R received the Spanish Researchers in the United Kingdom Society (SRUK/CERU) Winter Studentship to host A.M-A.

## Notes

### Competing Interest Statement

The authors have declared no competing interest.

