## Supporting Information for "Bi-functional particles for real-time acidification and proteolysis multiplex assay in macrophages"

**CONTENT:**

**Figure S1. Bi-functional particles in RAW 264.7 cells.**

**Figure S2. Real-time multiplex phagosome acidification and proteolysis analysis in RAW 264.7 cells.**

**Figure S3. Quality control of bi-functional particles.**

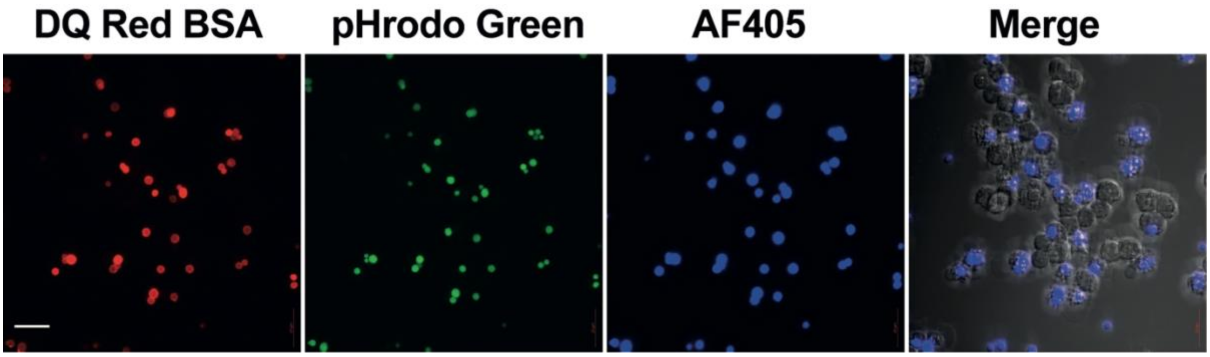


**Figure S1. Bi-functional particles in RAW 264.7 cells.** Immunofluorescence microscopy was performed in RAW 264.7 cells after 6 h post-uptake of bi-functional particles. DAPI in blue, pHrodo green in green, DQ red BSA in red, electronically switchable illumination and detection image (ESID) in white. Representative images are shown. Scale bar represents 10 µm.

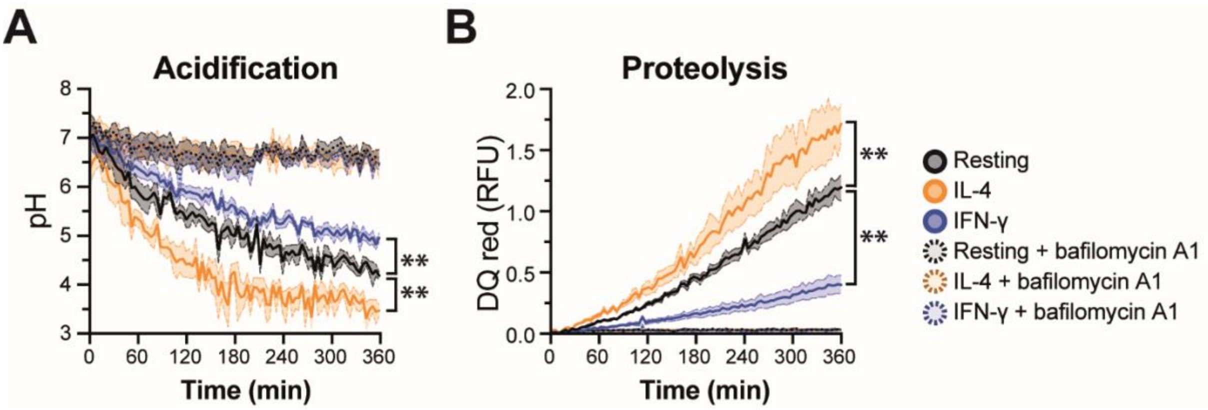


**Figure S2. Real-time multiplex phagosome acidification and proteolysis analysis in RAW 264.7 cells. A)** Acidification and **B)** proteolysis were measured in BMDMs untreated (resting) or treated with 20 µg/mL IFN-γ or 20 µg/mL IL-4 for 24 h. Bafilomycin A1 was used as a negative control of phagosome maturation. Friedman one-way ANOVA test followed by Dunn post hoc test. The statistical significance of the comparisons with resting is indicated as follows: **, $P \leq 0.01$. Error bars represent SEM of six biological replicates.

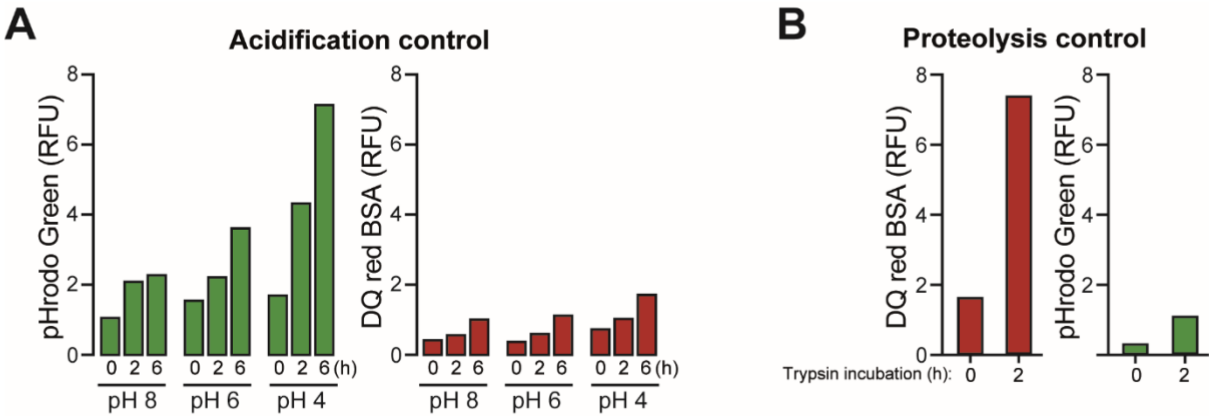


**Figure S3. Quality control of bi-functional particles. A**) Differet pH buffers for 6 h increased relative fluorescence units (RFU) in pHrodo green (490/530 nm) but not red fluorescence (585/625 nm). **B)** Bi-functional particles incubated for 2 h at 37°C with 1 μg/μL trypsin increased DQ red BSA fluorescence (585/625 nm), but not green fluorescence (490/530 nm).
